# A cell surface arabinogalactan-peptide influences root hair cell fate

**DOI:** 10.1101/787143

**Authors:** Cecilia Borassi, Javier Gloazzo Dorosz, Martiniano M. Ricardi, Laercio Pol Fachin, Mariana Carignani Sardoy, Eliana Marzol, Silvina Mangano, Diana Rosa Rodríguez Garcia, Javier Martínez Pacheco, Yossmayer del Carmen Rondón Guerrero, Silvia M. Velasquez, Bianca Villavicencio, Marina Ciancia, Georg Seifert, Hugo Verli, José M. Estevez

## Abstract

Root hairs (RHs) develop from specialized epidermal cells called trichoblasts, whereas epidermal cells that lack RHs are known as atrichoblasts. The mechanism controlling root epidermal cell fate is only partially understood. Root epidermis cell fate is regulated by a transcription factor complex that promotes the expression of the homeodomain protein GLABRA 2 (GL2), which blocks RH development by inhibiting ROOT HAIR DEFECTIVE 6 (RHD6). Suppression of GL2 expression activates RHD6, a series of downstream TFs including ROOT HAIR DEFECTIVE 6 LIKE-4 (RSL4 [Yi et al. 2010]) and their target genes, and causes epidermal cells to develop into RHs. Brassinosteroids (BRs) influence root epidermis cell fate. In the absence of BRs, phosphorylated BIN2 (a Type-II GSK3-like kinase) inhibits a protein complex that directly downregulates GL2 [Chen et al. 2014]. Here, we show that the genetic and pharmacological perturbation of the arabinogalactan peptide (AG) AGP21 in *Arabidopsis thaliana*, triggers aberrant RH development, similar to that observed in plants with defective BR signaling. We reveal that an *O*-glycosylated AGP21 peptide, which is positively regulated by BZR1, a transcription factor activated by BR signaling, affects RH cell fate by altering *GL2* expression in a BIN2-dependent manner. These results suggest that perturbation of a cell surface AGP disrupts BR responses and inhibits the downstream effect of BIN2 on the RH repressor GL2 in root epidermal cells. In addition, AGP21 also acts in a BR-independent, AGP-dependent mode that together with BIN2 signalling cascade controls RH cell fate.

**Significance:** In the plant *Arabidopsis thaliana*, the root epidermis forms in an alternating pattern atrichoblasts with trichoblast cells that end up developing root hairs (RHs). Atrichoblast cell fate is directly promoted by the transcription factor GLABRA2 (GL2) while the lack of GL2 allows RH formation. The loss of AGP21 peptide triggers an abnormal RH cell fate in two contiguous cells in a similar manner as brassinosteroid (BRs) mutants. In the absence of BR signaling, BIN2 (a GSK3 like-kinase) in a phosphorylated state, downregulate GL2 expression to trigger RH cell fate. The absence of AGP21 is able to repress *GL2* expression and activates the expression of RSL4 and EXP7 root hair proteins.

## Introduction

Plant roots not only anchor the plant into the soil but also allow them to absorb water and nutrients from the soil. Root hairs (RHs) are single cell protrusions developed from the epidermis that increase the root surface area exposed to the soil enhancing water and nutrients uptake. Many factors determine whether, or not, an epidermal cell will develop into a RH. These factors include both, environmental cues (such as nutrients in the soil) and signals from the plant itself, such as hormones like brassinosteroids (BRs), ABA, ethylene and auxin (Van Hengel et al. 2004; Masucci and Schiefelbein 1994, 1996; Kuppusamy et al., 2009). RH cell fate in the model plant *Arabidopsis* is controlled by a well-known developmental program, regulated by a complex of transcription factors composed by WEREWOLF (WER)-GLABRA3 (GL3)/ENHANCER OF GLABRA3 (EGL3)-TRANSPARENT GLABRA1 (TTG1) that promotes the expression of the homeodomain protein GLABRA 2 (GL2) (Ryu et al. 2005; Song et al. 2011; Schiefelbein et al. 2014; Balcerowicz et al. 2015), which ultimately blocks the root hair pathway by inhibiting ROOT HAIR DEFECTIVE 6 (RHD6) (Lin et al. 2015). The suppression of GL2 expression triggers epidermal cells to enter into the root hair cell fate program by the concomitant activation of RHD6 and a well-defined downstream gene network. As a consequence, RH and non-RH cell files are patterned alternately in rows within the root epidermis. In trichoblasts, a second transcription factor complex composed by CAPRICE (CPC)-GL3/EGL3-TTG1 suppresses GL2 expression (Schiefelbein et al. 2014), forcing cells to enter the RH cell fate program via concomitant RHD6 activation and downstream TFs, including RSL4, and RH genes (Yi et al. 2010). The plant steroid hormones, BRs play essential roles in regulating many developmental processes (Savaldi-Goldstein et al., 2007; 2010; Hacham et al., 2011; Yang et al., 2011). BRs are perceived by the receptor kinase BRASSINOSTEROID INSENSITIVE 1 (BRI1) (Li & Chory, 1997; Hothorn et al., 2011; She et al., 2011). One of the BRI1 substrate, BR-SIGNALING KINASE (BSK), transduces the BR signaling through *bri1* SUPPRESSORS 1 (BSU1) to inactivate a GSK3-like kinase BRASSINOSTEROID INSENSITIVE 2 (BIN2), which triggers high levels of the dephosphorylated form of transcriptional factors BRI1 EMS SUPPRESSOR 1 (BES1)/BRASSINAZOLE RESISTANT 1 (BZR1) in the nucleus to regulate gene expression (Yan et al. 2009; Yang et al., 2011). In recent years, a molecular mechanism was proposed by which BR signaling controls RH cell fate by inhibiting BIN2 phosphorylation activity to modulate GL2 expression (Chen et al. 2014). In atrichoblasts, BIN2 phosphorylates TTG1, controlling protein complex TTG1-WER-GL3/EGL3 activity, and stimulating *GL2* expression (Chen et al. 2014).

Plant cell surface proteoglycans known as arabinogalactan proteins (AGPs) function in a broad developmental processes such as cell proliferation, cell expansion, organ extension, and somatic embryogenesis (Tan et al. 2004; Seifert & Roberts 2007; Pereira et al. 2015; Ma et al. 2018). The precise mechanisms underlying AGP action in these multiple processes are completely unknown (Ma et al. 2018). AGP peptides are post-translationally modified in the ER-Golgi, undergoing signal peptide (SP) removal, proline-hydroxylation/Hyp-*O*-glycosylation, and C-terminal GPI anchor signal (GPI-AS) addition (Schultz et al. 2004; Ma et al. 2018). Processed mature AGP-peptides are 10–13 amino acids long and bear few putative *O*-glycosylation sites (*O*-AG). Few prolines in the AGP peptides are hydroxylated in vivo as Hyp (Hyp=O), suggesting that AGP peptides are *O*-glycosylated at maturity (Schultz et al. 2004). All these posttranslational modifications make the study of AGPs very complex with almost no defined biological functions for any individual AGP (Ma et al. 2018). Interestedly, in this work we have identified that disruption of plant specific AGPs, and in particular of a single *O*-glycosylated AGP peptide (AGP21), interfere in a specific manner with BR responses and BIN2 downstream effect on the repression of RH development. We have found that an *O*-glycosylated AGP21-peptide positively regulated by the BR transcription factor BZR1, impacts on RH cell fate in a BIN2-dependent manner by controlling GL2 expression.

## Results and Discussion

### AGP perturbation influences root hair (RH) cell fate programming

To determine whether *O*-glycosylated AGPs regulate specific RH developmental processes, we exposed roots of *Arabidopsis thaliana* to β-glucosyl Yariv (β-Glc-Y), which specifically binds structures in the *O*-glycans of AGPs: oligosaccharides with at least 5–7 units of 3-linked *O*-galactoses (Yariv et al. 1967; Kitazawa et al. 2013). β-Glc-Y–linked AGP complexes on the cell surface induce AGP aggregation and disrupt native protein distribution, triggering developmental reprogramming (Guan & Nothnagel 2004; Sardar et al. 2006). α-mannosyl Yariv (α-Man-Y), an analogue that does not bind to AGPs, served as the control. While α-Man-Y treatment did not affect RH cell fate (≈2–5% of total RHs that are contiguous), β-Glc-Y treatment increased contiguous RH development (≈30-35%) (**Figure S1A**), suggesting that *O*-glycosylated AGPs influence RH cell fate.

To test whether *O*-glycans on hydroxyproline-rich glycoproteins (HRGPs) alter RH cell fate, we blocked proline 4-hydroxylase enzymes (P4Hs) that catalyse proline (Pro)-hydroxylation into hydroxyl-proline units (Hyp), the subsequent step of HRGP *O*-glycosylation (Velasquez et al. 2011, 2015a). Two P4H inhibitors, α,α-dipyridyl (DP) and ethyl-3,4-dihydroxybenzoate (EDHB), prevent Pro-hydroxylation (Barnett 1970; Majamaa et al. 1986); both increased contiguous RH development to ≈15–20% (**Figure S1B**). Additionally, *p4h5* (a key P4H in roots [Velasquez et al. 2011; 2015a]) and four glycosyltransferase (GT) mutants defective in AGP and related proteins *O*-glycosylation (*hpgt* triple mutant; *ray1*, *galt29A*, and *fut4 fut6*) (see **Table S1**) showed significantly increased (≈8–20%) ectopic RH development (**Figure 1A**), substantiating the previous report that the triple mutant *hpgt* mutant has an increased RH density (Ogawa-Ohnishi & Matsubayashi 2015). These mutants were mostly insensitive to β-Glc-Y; however, the treatment increased the number of contiguous RHs in *fut4 fut6*, although to a lesser extent than in the wild type (**Figure 1B**). β-Glc-Y inhibits root cell expansion (Willats & Knox 1996; Ding & Zhu 1997). Glycosyltransferase (GT) mutations affecting extensin (EXTs) and related proteins *O*-glycosylation (e.g. *rra3* and *sgt1 rra3*; **Table S1**) drastically affect RH cell elongation (Velasquez et al. 2015b). Intriguingly, these mutations did not affect RH cell fate, and β-Glc-Y stimulated ectopic RH development as in Wt Col-0, indicating that EXT *O*-glycosylation might not function in RH cell fate reprogramming (**Table S1**, **Figure 1C**), and specifically *O*-glycans attached to AGPs and related glycoproteins do. *P4H5* and *AGP-related GTs (e.g. RAY1, GALT29A, HPGT1-HPGT3 and FUT4/FUT6)*, are expressed in the root epidermis elongation and differentiation zones (**Figure S2**). Under-arabinosylated AGPs in *ray1* and under–*O*-fucosylated AGPs in *fut4 fut6* show similar root growth inhibition (Liang et al. 2013; Trypona et al. 2014), highlighting a key role for AGP *O*-glycans in regulating root cell development, albeit by unknown mechanisms. These results using DP/EDHB and β-Glc-Y treatments as well as mutants in the AGPs *O*-glycosylation pathway suggest that AGPs and related proteins might be involved in RH cell fate.

**Figure 1.**
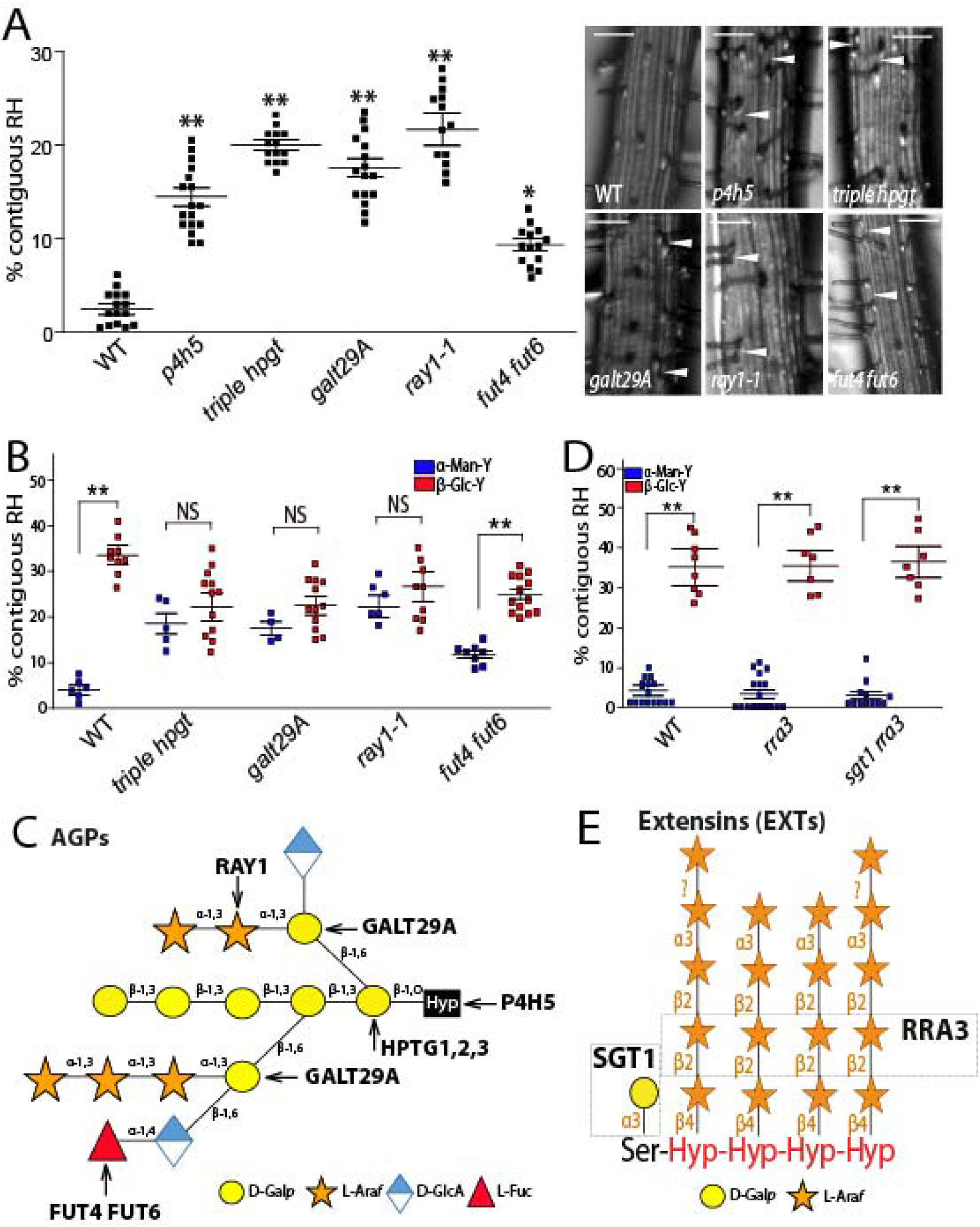
Contiguous RH phenotype in *O*-underglycosylated AGPs phenocopy BR mutants. (A) RH phenotype in the *p4h5* mutant and in four glycosyltransferase mutants (*triple hpgt, ray1, galt29A*, and *fut4 fut6*) that act specifically on AGP *O*-glycosylation. Right, selected pictures. Arrowheads indicated two contiguous RHs. Scale bar= 50 μm. (B) RH phenotype in three glycosyltransferase (GT) mutants (*triple hpgt, ray1, galt29A* and *fut4 fut6*) that act specifically on AGP *O*-glycosylation. Effect on contiguous RH phenotype in roots treated with 5μM α-Mannosyl Yariv (α-Man-Y) or 5μM β-Glucosyl Yariv (β-Glc-Y). (C) The mutants used in (B) for the GTs involved in AGP *O*-glycosylation are indicated. (D) RH phenotype in two glycosyltransferase mutants (*rra3* and *rra3 sgt1*) that act specifically on EXT *O*-glycosylation. Effect on contiguous RH phenotype in roots treated with 5μM α-Mannosyl Yariv (α-Man-Y) or β-Glucosyl Yariv (β-Glc-Y). (E) The mutants used in (D) for the GTs involved in EXT *O*-glycosylation are indicated. (A, B and D) *P*-value of one-way ANOVA, (**) P<0.001, (*) P< 1. NS= not significant different. Error bars indicate ±SD from biological replicates. See also Figure S1-S4.

### The AG peptide AGP21 influences RH cell fate

Brassinosteroid (BR) signaling regulates RH cell patterning (Cheng et al. 2015). The BR-insensitive mutant, *bri1-116*, and *bak1* developed many (≈20%-25%) contiguous RH cells (**Figure S3A**), resembling plants subjected to β-Glc-Y and DP/EDHB treatments (**Figure S1**). *p4h5*, *hpgt* triple *mutant, ray1-1, galt29A*, and *fut4 fut6* mutants exhibited similar phenotypes, suggesting that interplay between cell surface AGPs and BR signaling determines RH cell fate. As chromatin-immunoprecipitation (ChIP)-sequencing and RNA-sequencing indicate that BZR1 directly upregulates *AGP* expression, most predominantly *AGP21* (Sun et al. 2010), we investigated how root epidermal BR signalling regulates *AGP21* expression. Since the *AGP21* regulatory region contains one BZR1 binding motif (E-BOX, CATGTG at −279 bp relative to ATG start codon), we tested whether BR directly modulates *AGP21* expression. Compared with no treatment, 100 nM BL (brassinolide, BR’s most active form) enhanced of both AGP21p∷GFP (transcriptional reporter) and AGP21p∷V-AGP21 (V= Venus tag; translational reporter) expression (**Figure S3B–C**). Expression of *AGP21p*∷*GFP* in *bri1-116* resulted in lower *AGP21* signal than in untreated wild type (**Figure S3B**), confirming that BR-mediated BZR1 controls *AGP21* expression in the root. Trichoblasts and atrichoblasts expressed V-AGP21 peptide in a discontinuous pattern (**Figure S1C**), indicating that some root epidermal cells expressed AGP21 but some lacked it. Treatment with β-Glc-Y—but not α-Man-Y—resulted in excess AGP21p∷Venus-AGP21 at transverse cell walls (**Figure S1C**) confirming the expect effect on aggregating AGPs at the cell surface. These results might indicate AGP21 as a possible link between RH cell fate phenotype and BR responses in root epidermal cells.

Although we screen for abnormal RH cell fate in several AGP-peptide mutants, only AGP21 deficient mutant *agp21* (**Figure S4A–B**), exhibited ectopic contiguous RHs at high levels (≈20%) (**Figure 2B**). Both *AGP21* expression under its endogenous promoter (*AGP21p∷V-AGP21/agp21*) and overexpression (*35Sp∷V-AGP21/agp21*) restored a wild type RH phenotype and patterning to *agp21* (**Figure 2B**), confirming that deficient *AGP21* expression causes contiguous RH development. Furthermore, while β-Glc-Y treatment triggered up to ≈35% of contiguous RH (vs. ≈2–5% induced by α-Man-Y) in the wild type (**Figure S1A**), it induced no additional anomalous RH in *agp21* (vs. α-Man-Y treatment or untreated roots) (**Figure 2B**). We tested whether the closely related BZR1-induced peptide AGP15 functions with AGP21 (Sun et al. 2010). *agp15* (**Figure S4C–D**) exhibited a milder phenotype than *agp21*, and the double *agp15 agp21* double mutant had no additional effects to *agp21* (**Figure S4E**). Together, these results confirm that β-Glc-Y might affect *O*-glycosylated AGP21 to stimulate contiguous RH development.

**Figure 2.**
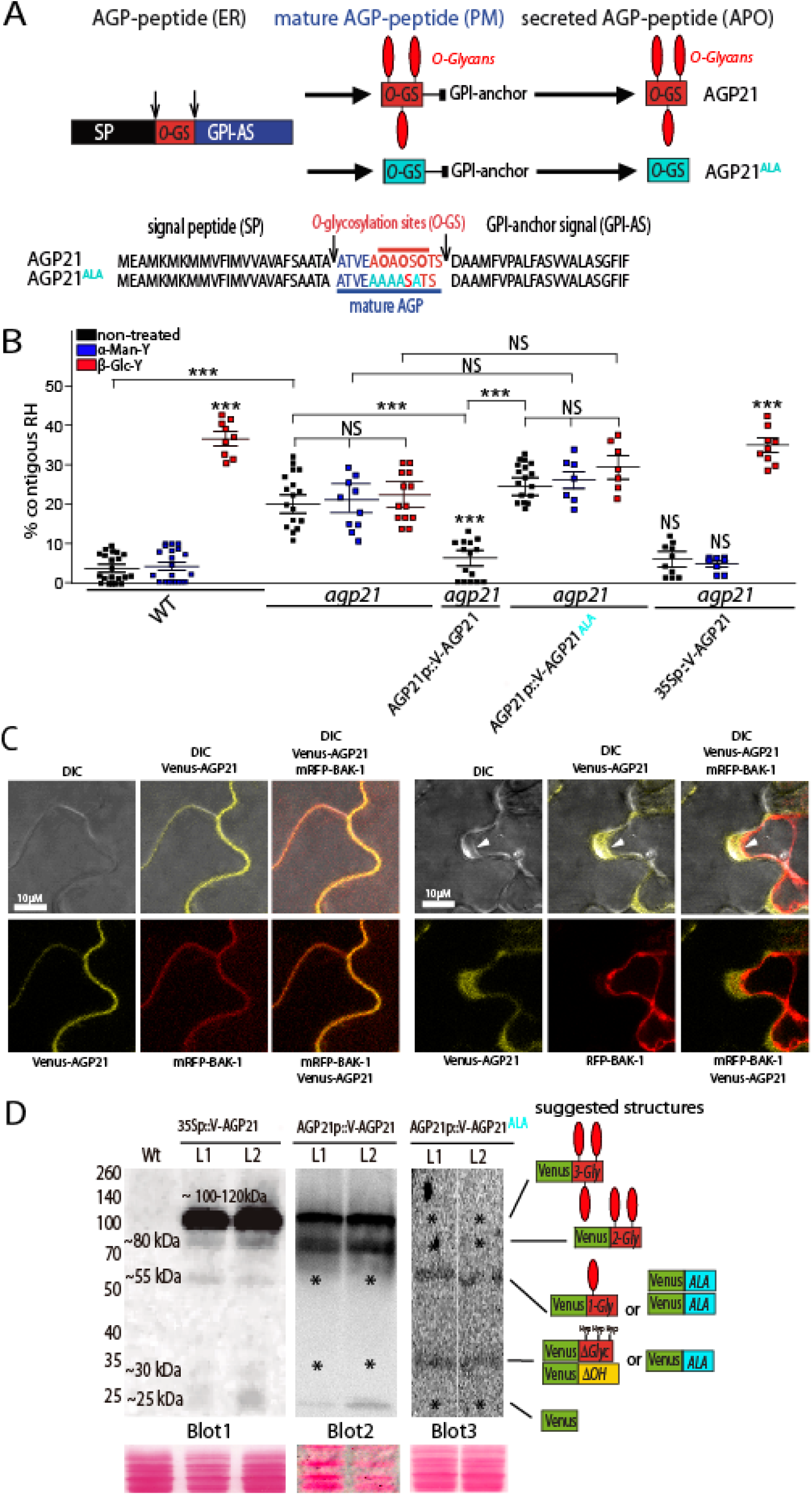
*O*-glycosylated AGP21 peptide at the cell surface modulates RH cell fate. (A) Identified AGP21 peptide acting on root epidermis development. AGP21 peptide sequence and its posttranslational modifications carried out in the secretory pathway. The mature AGP21 peptide contains only 10-13 aa in length. APO= Apoplast. ER=Endoplasmic Reticulum. GPI anchor= GlycosylPhosphatidylInositol (GPI) anchor. PM=Plasma membrane. (B) Contiguous RH phenotype in *agp21*, complemented *agp21* mutant with AGP21p∷V-AGP21 and with 35Sp∷V-AGP21 constructs as well as AGP21p∷V-AGP21^ALA^ expression in *agp21*. Only one line is shown. *P*-value of one-way ANOVA, (**) P<.001, (*) P<.01. NS= not significant differences. Error bars indicate ±SD from biological replicates. (C) Co-localization of AGP21-Venus with BAK1-mRFP at the plasma membrane of epidermal cells in *Nicothiana Benthamiana*. Scale bar= 10 μm. Cross section of expression levels across BAK1-RFP coexpressed with AGP21-Venus. On the left, plasmolysis was induced with 800 mM Mannitol uncovering an apoplastic plus plasma membrane AGP21 localization. Scale bar= 10 μm. Arrowheads indicate plasma membrane located AGP21. Scale bar= 50 μm. (D) Immunoblot analysis of two stable lines expressing 35Sp∷V-AGP21 (L1-L2) and two lines expressing AGP21p∷V-AGP21 (L1-L2) and two lines expressing AGP21p∷V-AGP21^ALA^ (L1-L2). Each blot is an independent experiment. Putative Venus-AGP21 structures are indicated on the right based on the apparent molecular weight. *O*-glycans are indicated as red elongated balloons. ΔOH = non-hydroxylated. ΔGly = without *O*-glycans. 1-Gly to 3-Gly = 1 to 3 sites with Hyp-*O*-glycosylation. Asterisk indicates missing AGP21 glycoforms or lack of Venus protein. See also Figure S4-S6.

### *O*-glycosylation is required for the correct targeting of the AGP21 peptide to the plasma membrane-apoplastic space

To determine whether functional AGP21 requires *O*-glycosylation, three putative *O*-glycosylation sites were mutated (Pro→Ala) (**Figure 2A**) and driven by the endogenous *AGP21* promoter in *agp21* (*AGP21p∷V-AGP21^ALA^/agp21*). Mass spectrometry had detected that all three proline units (Pro/P) within the AGP21 sequence ATVEAPAPSPTS can be hydroxylated as ATVEA<u>O</u>A<u>O</u>S<u>O</u>TS (Hyp= <u>O</u>) (Schultz et al. 2004), indicating likely sites for *O*-glycosylation. Even though AGP21^ALA^ protein was detected in root epidermal cells (**Figure S5B**), AGP21^ALA^ failed to rescue the agp21 RH phenotype (**Figure 2B–C**), suggesting that Hyp-linked *O*-glycans in AGP21 are required for its function in RH cell fate. Moreover, β-Glc-Y treatment did not induce anomalous RH cell fate in AGP21^ALA^ plants. Then, we examined whether AGP21 expressed in Nicotiana benthamiana colocalized with the BRI1 co-receptor BAK1 (**Figure 2C**). V-AGP21 partially colocalized with BAK1-mRFP protein (**Figure 2C**). When epidermal cells were plasmolyzed, most AGP21 signal localized to the apoplast but some remained close to the PM (**Figure S8B**). V-AGP21^ALA^, however, never reached the cell surface; retention in the secretory pathway could indicate that *O*-glycans direct AGP to the PM–cell surface (**Figure S5A–B**). These data is in agreement with previous reports of a requirement for *O*-glycans in the secretion and targeting of AGPs and related fasciclin-like AGPs (Xu et al 2008; Xue et al 2017).

We tested the hypothesis that AGP21 is processed and modified during its synthesis along the secretory pathway. Using immunoblot analysis, we examined the apparent molecular weight of AGP21 peptide in transient AGP21-overexpressing plants and in *AGP21p∷V-AGP21 plants* (**Figure 2D**). In the overexpressing plants, most AGP21 peptide was detected as a strong broad band around ≈100–120 kDa with minor bands at ≈80 and ≈55 kDa, whereas endogenously driven AGP21 produced a stronger band at ≈80 kDa and lacked the band at ≈55 kDa, suggesting that, in both cases, AGP21 peptide might be present in a putative tri-*O*-glycosylated form. Mature peptide with no posttranslational modifications is approximately 30 kDa; the extra bands could be interpreted as intermediate single- and di-*O*-glycosylated forms of AGP21 peptide. An apparent molecular shift of ≈25–30 kDa for each putative *O*-glycosylation site in AGP21 accords with AGP14 peptide, whose protein sequence is highly similar (Ogawa-Ohnishi & Matsubayashi 2015), and with the electrophoretic migration of an AGP-xylogen molecule that contains two arabino-galactan-*O*-Hyp sites (Motose et al. 2004). V-AGP21^ALA^, which lacks *O*-glycans, is not targeted to the cell surface, formed puncta structures (**Figure S5B**) and showed one band close to ~55 kDa (**Figure 2D**) and one band close to ~30 kDa. It is hypostatized here that lack of *O*-glycans V-AGP21^ALA^’s may cause to self interactions and this is compatible with the punctuated structure visualized in the root epidermal cells (**Figure S5B**). A detailed analysis is required to characterize *O*-glycosylation in AGP21 peptide although it is technically challenging due to its carbohydrate complexity.

### *O*-glycans stabilize AGP21 peptide’s functional conformation

To address the effect of *O*-glycan on the conformation and stability of AGP21 peptide, we modeled a minimal, 15-sugar Hyp-*O*-linked arabinogalactan (AG) structure ([ATVEAP(O)AP(O)SP(O)TS], **Figure S6A–B**). This is the simplest carbohydrate structure characterized for a single AGP synthetic peptide (Tan et al. 2004), although more complex structures were described for several AGPs (Kitazawa et al.2013). To assess the conformation of AGP21 peptide and the effect of *O*-glycosylation, molecular dynamics (MD) simulations considered three non-glycosylated peptides (with alanines [nG-Ala], prolines [nG-Pro], or hydroxyprolines residues [nG-Hyp], respectively) and one *O*-glycosylated peptide with three Hyp-*O*-glycans (**Figure S6C**). In the MD simulations, the root mean square deviation (RMSD) varied up to ≈6 Å (**Figure S6D**), indicating that peptide structure may have deviated from the starting type-II polyproline helix. By contrast, larger conformational stabilization effects were observed in the *O*-glycosylated peptide (**Figure S6E**). Individual residue RMSF analysis indicated that the peptide’s stiffer region depended on the MD conditions applied (**Figure S6F**). To characterize conformational profiles, we measured the angle formed by four consecutive alpha carbon atoms (ζ angle) (**Table S3**). The ζ angle of a type-II polyproline helix is −110 ± 15°. In this context, the *O*-glycosylated AOAOSOTS peptide structure is slightly extended between Pro2–Thr7, as observed by ζ angles 2–4 closer to 180° (**Table S3**). Our analysis suggests that *O*-linked glycans affect the conformation and stability of AGP21 peptide. How this conformational change in mature AGP21 peptide without *O*-glycans affects its function in RH cell determination remains unclear.

### AGP21 acts in a BIN2-dependent pathway to define RH cell fate

We hypothesized that disrupting AGPs activity with β-Glc-Y, a lack of AGP21 peptide (*agp21*), or abnormal glycosylation on AGP and related proteins, would interfere with BR responsiveness and RH cell fate. We treated the triple mutant gsk (*gsk triple: bin2-3 bil1 bil2*; BIL1, BIN2-like 1 and BIL2, BIN2-like 2), which almost completely lacks RH cells [1], with 5 μM β-Glc-Y treatment. Gsk triple exhibited few contiguous RH cells before and after the treatment (**Figure 3**), suggesting that β-Glc-Y requires BIN2-BIL1-BIL2 to alter RH cell fate. Interestingly, β-Glc-Y induced ≈40-45% contiguous RHs (**Figure 3**) in the constitutively active mutant *bin2-1* (Li & Nam 2002). These data suggest that the AGP-mediated RH cell fate reprogramming requires active BIN2, BIL1, and BIL2 proteins (**Figure 3A**).

**Figure 3.**
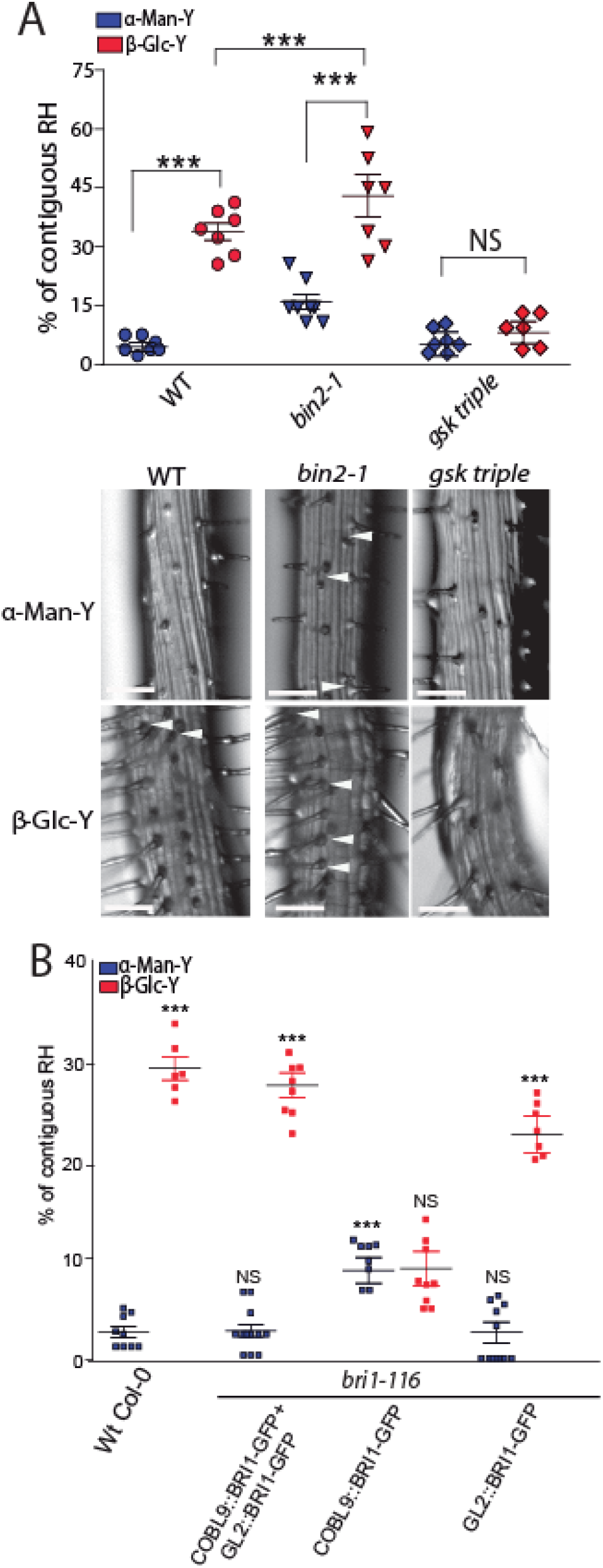
Perturbation of AGPs requires active BRI1 expression in atrichoblast cells and downstream BIN2-BIL1-BIL2 proteins to triggers changes in RH cell fate. (A) Contiguous RH phenotype in roots treated with 5μM β-Glucosyl Yariv (β-Glc-Y) or 5μM α-Mannosyl Yariv (α-Man-Y). Scale bar= 20 μm. *P*-value of one-way ANOVA, (***) P<0.001, (*) P<0.05. NS= not significant differences. Error bars indicate ±SD from biological replicates. Arrowheads indicated two contiguous RHs. (B) Effect of the BRI1 differential expression on the development of contiguous RH. BRI1 is active when expressed in atrichoblast cells (under GL2 promoter). See also Figure S5.

As *BRI1* expression is similar in trichoblasts and atrichoblasts (Fridman et al., 2014), we sought to determine whether BRI1 and downstream BR responses act differently in these cell types during RH cell fate determination (**Figures 3B**). We examined the effect of cell type-specific *BRI1* expression on the percentage of contiguous RHs in three plant lines expressing BRI1-GFP, all in the *bri1-116* background: trichoblast-only (*COBL9p∷BRI1-GFP/bri1-116*), atrichoblast-only (*GL2p∷BRI1-GFP/bri1-116*), and expression in both cell types (*GL2p∷BRI1-GFP + COBL9p∷BRI1-GFP/bri1-116*) (Hacham et al., 2011; Fridman et al., 2014). BRI1 expression in atrichoblasts only did not rescue *bri1-116* (plants showed abundant contiguous RHs), the line that expressed BRI1 in trichoblasts or in both cell types were similar to wild type (**Figure 3B**). Additionally, only *COBL9p∷BRI1/bri1-116* where BRI1 is missing only in atrichoblast cells, was completely insensitive to β-Glc-Y while the other two lines exhibited more contiguous RHs. These data may imply that only the BR-BRI1 pathway in atrichoblasts is active to promote ectopic RH development and is also sensitive to AGP disruption.

### Disturbance or absence of AGP21 blocks *GL2* expression

We tracked epidermal cell fate and analyzed β-Glc-Y and α-Man-Y’s translational effects on several markers: an early RH marker (RHD6p∷RHD6-GFP), a downstream transcription factor (RSL4p∷RSL4-GFP), a late RH marker (EXP7p∷EXP7-GFP), and an atrichoblast marker GL2 (GL2p∷GL2-GFP) (**Figure 4A–D**). β-Glc-Y, not α-Man-Y, repressed GL2 expression and enhanced RHD6, RSL4 and EXP7 expression in contiguous epidermal cells (**Figure 4A–E**). This corroborates the effects of both β-Glc-Y and deficiencies in the AGP *O*-glycosylation pathway on contiguous epidermis cell development. Then, when we expressed RSL4p∷RSL4-GFP in *agp21*, two contiguous epidermis cells showed GFP expression, while this rarely occurred in wild type roots. The transcriptional reporter GL2p∷GFP/*agp21* showed discontinuous RH patterning similar to β-Glc-Y treatment (**Figure 4B** and **4D**). This result implies feedback between the lack of AGP21, GL2 repression, and RHD6-RSL4 and EXP7 upregulation in contiguous epidermal cell development (**Figure 4E**). Constitutively active *bin2-1* phenocopies *agp21* and β-Glc-Y treatment: it represses GL2 expression in some epidermal cells and enhances EXP7-GFP in contiguous epidermal cells, stimulating contiguous RH development (**Figure 4F–G**). To test whether AGP21 (and AGPs in general), affect BR responses, we treated roots with 100 nM BL. Wild type roots exhibited repressed RH development as previously reported (Cheng et al. 2014); *agp21* and three GT mutants (*triple hpgt, ray1* and *galt29A*) defective in AGP *O*-glycosylation (**Table S1**) were unaffected by BL treatment (**Figure S6C**), suggesting that *O*-glycosylated AGP21 (and AGPs) are required for promoting BR responses and downstream signalling on RH cell fate.

**Figure 4.**
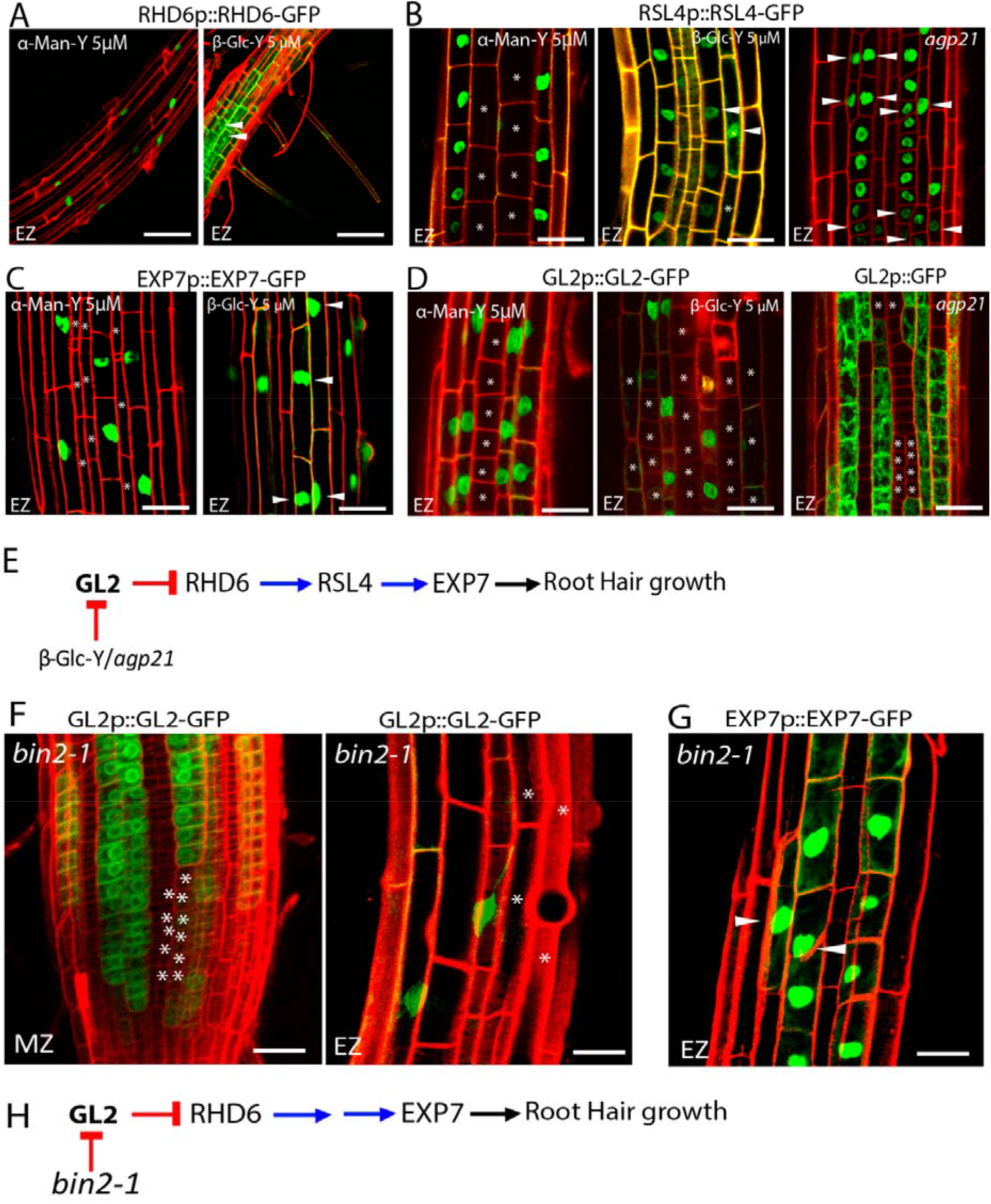
AGPs disruption, the lack of AGP21, and *bin2-1* block the RH repressor GLABRA2 (GL2) and triggers RHD6-RSL4-EXP7 expression in some atrichoblast cells. The effect of β-Glucosyl Yariv (β-Glc-Y), α-Mannosyl Yariv (α-Man-Y), and the absence of AGP21-M-M peptide were monitored on several markers to study epidermis cell fate. (A) RHD6 (RHD6p∷RHD6-GFP) as an early RH marker. (B) A downstream RHD6 factor RSL4 (RSL4p∷RSL4-GFP). (C) The RSL4-gene target EXP7 (EXP7p∷EXP7-GFP). (D) The main RH repressor GL2 (GL2p∷GL2-GFP). (A-D) Arrowheads indicate expression of a given marker in two contiguous epidermis cell lines. Asterisks indicate absence of expression. Scale bar= 20 μm. (E) Proposed sequence of events triggered by β-Glucosyl Yariv (β-Glc-Y) or the lack of AGP21 peptide that leads to abnormal RH development. (F) GL2 expression in the *bin2-1* background in the Meristematic Zone (MZ) and Elongation Zone (EZ) of the root. (G) The RH marker EXP7 expression in the *bin2-1* background in the Elongation Zone (EZ) of the root. (F-G) Arrowheads indicate expression of a given marker in two contiguous epidermal cell lines. Asterisks indicated absence of expression. Scale bar= 10 μm. (H) Proposed sequence of events triggered by *bin2-1* that leads to abnormal RH development. See also Figure S6.

## Conclusions

In root epidermal cells, atrichoblast fate is the default, while environmental as well as endogenous cues like high levels of BRs promotes *GL2* expression in atrichoblasts to repress RH development (Cheng et al. 2014). In the absence of BRs, active P-BIN2 represses *GL2* expression and *RHD6* and *RSL4* expression proceeds, triggering RH development in atrichoblasts and producing contiguous RHs. Perturbed AGPs and the lack of AGP21 peptide at the cell surface stimulate ectopic RH development similar to that observed in BR mutants. BZR1 regulates *AGP21* expression and the *O*-glycosylated cell surface peptide AGP21 modulates RH cell fate. We propose a model, in which the *O*-glycosylated AGP21 peptide and BR responses are both dependent on BIN2 (and BIL1-BIL2)-mediated responses, controlling RH cell fate (**Figure S7**). It still unclear how the cell surface peptide AGP21 is able to trigger a change in RH cell fate in a BIN2-dependent manner. One possibility is that AGP21 peptide might modify the responsiveness to BRs of the co-receptors BRI1-BAK1. In line with this, we failed to detect a direct interaction between V-AGP21 and BAK1-mRFP in a transient expression system (results not shown). Nonetheless, measuring direct physical interactions between *O*-glycosylated AGP21 and BRI1–BAK1 proteins in the apoplast–PM space is a challenge for a future study. In concordance with this scenario, other GPI anchor proteins (e.g. like LORELEI-like-GPI-anchored protein 2 and 3, LRE/LLG2,3) are able to interact with CrRLK1s (e.g. FERONIA and BUP1,2/ANXUR1,2) in the cell surface of polar growing plant cells (Li et al. 2015; 2016; Lui et al. 2016; Ge et al. 2019; Feng et al.2019). These results imply an interesting parallel between plant AGPs and animal heparin sulfate proteoglycans (HSPGs), which are important co-receptors in signaling pathways mediated by growth factors, including members of Wnt/Wingless, Hedgehog, transforming growth factor-β, and fibroblast growth factor family members (Lin 2004). A second scenario is that AGP21 peptide and BR co-receptors BRI1-BAK1 do not interact in the cell surface and both influence by different pathways BIN2 activity and the downstream RH cell fate program. If this is the case, AGP21 may require others proteins to transduce the signal toward BIN2 in the cytoplasm. Future work should investigate which of these two hypotheses might explain the role of AGP21 peptide in RH cell fate.

## Materials and Methods

### Growth conditions

All plant materials used in this study were in the Columbia-0 ecotype background of *Arabidopsis thaliana*. Seeds were sterilized and placed on half-strength (0.5X) Murashige and Skoog (MS) medium (Sigma-Aldrich) pH 5.8 supplemented with 0.8% agar. For root measurements, RNA extraction and confocal microscopy 7-day old seedlings were grown on square plates placed vertically at 22°C with continuous light, after stratification in dark at 4°C for 5 days on the plates. Seedlings on plates were transferred to soil and kept in the greenhouse in long-day conditions to obtain mature plants for transformation, genetic crossing, and amplification of seeds.

### Plant material

For identification of homozygous T-DNA knockout lines, genomic DNA was extracted from rosette leaves. Confirmation by PCR of a unique band corresponding to T-DNA insertion in the target genes AGP15 (At5G11740: SALK_114736), AGP21 (At1G55330: SALK_140206), HPGT1-HPGT3 (AT5G53340: SALK_007547, AT4G32120: SALK_070368, AT2G25300: SALK_ 009405) GALT29A (At1G08280: SALK_030326; SALK_113255; SAIL_1259_C01) and RAY1 (At1G70630: SALK_053158) were performed using an insertion-specific LBb1.3 for SALK lines or Lb1 for SAIL lines. Primers used are listed in **Table S4**. The stable transgenic lines used in this study are summarized in **Table S2**.

### Pharmacological treatments

ethyl-3,4-dihydrohydroxybenzoate (EDHB) and α,α-Bipyridyl (DP) D216305 SIGMA-ALDRICH were used as P4Hs inhibitors. DP chelates the cofactor Fe^2+^ [9] and the EDHB interacts with the oxoglutarate-binding site of P4Hs (Majamaa et al. 1986). Specific Yariv phenylglycoside (for 1,3,5-tri-(p-glycosyloxyphenylazo)-2,4,6-trihydroxybenzene), β-glucosyl Yariv phenylglycoside (β-Glc-Yariv) was used for AGP-depletion (Kitazawa et al.2013). α-mannosyl Yariv phenylglycoside (α-Man-Yariv) was used as negative control for phenylglycoside treatment. Both, β-Glc-Y and α-Man-Y are Yariv-phenylglycosides and its specificity for AGPs relies on the β-configuration of the glycosyl residues attached to the phenylazotrihydroxybenzene core (Yariv et al. 1967). DP, EDHB, or Yariv reagents were added to MS media when MS plates were made. Seedlings were grown for 4 days in MS 0.5X media and then transferred for 3 days more to MS 0.5X plates with DP, EDHB, or Yariv reagents at the concentration indicated.

### Quantification of RH cell fate

In order to determine the RH patterning, images of root tips were taken using an Olympus stereomicroscope at maximum magnification (50X). The presence of contiguous RH was analyzed using ImageJ, starting from the differentiation zone to the elongation zone. The amount of contiguous RH was expressed as a percentage of total RH for rectangular root areas of 200 μm in width x 2mm in length (n=20) with three biological replicates. Quantitative and statistical analysis was carried on using GraphPad software. To analyze the alteration in RH cell fate, root cell walls of reporter lines were stained with 5 µg/ml propidium iodide and confocal microscopy images were taken using a Zeiss LSM 710 Pascal microscope, 40X objective N/A= 1.2.

### AGP21 variants

AGP21 promoter region (AGP21p) comprising 1,5 Kbp upstream of +1 site was amplified by PCR and cloned into pGWB4 to obtain AGP21p∷GFP construct. Synthetic DNA was designed containing full length AGP21 cDNA and Venus fluorescent protein cDNA between AGP21 signal sequence and the mature polypeptide (Venus-AGP21), containing GatewayTM (Life Technologies) attB1 and attB2 sites. Recombinase-mediated integration of the PCR fragment was made into pEntry4Dual. pEntry4Dual/Venus-AGP21 construction was recombined into the vector pGWB2 (Invitrogen, Hygromicyn R) in order to overexpress Venus-AGP21 under 35S mosaic virus promoter (35Sp∷Venus-AGP21). Also, Venus-AGP21 construct was cloned into pGWB1 (no promoter, no tag) and AGP21p was sub-cloned in the resulting vector to express AGP21 reporter under the control of its endogenous promoter (AGp21p∷Venus-AGP21). Wild type and T-DNA *agp21* mutant plants were transformed by using Agrobacterium (strain GV3101+pSoup). Plants were selected with hygromycin (30 μg/ml) and several independent transgenic plants were isolated for each construct. At least three homozygous independent transgenic lines of Col-0/AGP21p∷GFP, *agp21*/AGP21p∷Venus-AGP21 and *agp21*/35Sp∷AGP21-GFP were obtained and characterized.

### Gene expression analysis

For RT-PCR analysis, total RNA was isolated from roots of 7-day-old seedlings using RNeasy Plant Mini Kit (Qiagen) according to the manufacturer’s instructions. cDNA synthesis was achieved using M-MLV reverse transcriptase (Promega). PCR reactions were performed in a T-ADVANCED S96G (Biometra) using the following amplification program: 4 min at 95°C, followed by 35 cycles of 20 secs at 95°C, 30 secs at 57°C and 30 secs at 72°C. RT-PCR was performed to assess AGP15 and AGP21 transcript levels in wild type and T-DNA mutant *agp15* and *agp21*. PP2A was used as an internal standard. All primers used are listed in **Table S4**.

### Confocal microscopy

Confocal laser scanning microscopy was performed using Zeiss LSM 510 Meta and Zeiss LSM 710 Pascal. Fluorescence was analyzed by using laser lines of 488 nm for GFP or 514 nm for YFP excitation, and emitted fluorescence was recorded between 490 and 525 nm for GFP and between 530 and 600 nm for YFP (40X objective, N/A= 1.2). Z series was done with an optical slice of 2µm, and intensities was summed for quantification of fluorescence along a segmented line using plot profile command in Image J, five replicates for each of five roots were observed.

### AGP21 Immunoblotting detection

Proteins were extracted from roots of 7-day-old seedlings using extraction buffer (20mM TRIS-HCl pH8.8, 150mM NaCl, 1mM EDTA, 20% glycerol, 1mM PMSF, 1X protease inhibitor Complete^®^ Roche) at 4°C. After centrifugation at 21.000*g* at 4°C for 20min, protein concentration in the supernatant was measured and equal protein amounts were loaded onto a 6% SDS-PAGE gel. Proteins were separated by electrophoresis and transferred to nitrocellulose membranes. Anti-GFP mouse IgG (Roche Applied Science) was used at a dilution of 1:1.000 and it was visualized by incubation with goat anti-mouse IgG secondary antibodies conjugated to horseradish peroxidase (1:10.000) followed by a chemiluminescence reaction (Clarity ™ Western ECL Substrate, BIO-RAD).

### Transient expression assays in *Nicotiana benthamiana*

To test the sub-cellular localization of AGP21, 5-day-old *N. benthamiana* leaves were infiltrated with *Agrobacterium* strains (GV3101) carrying 35Sp∷Venus-AGP21 and BAK1-RFP constructs. After 2 days, images of the lower leaf epidermal cells were taken using a confocal microscope (LSM5 Pascal) to analyze Venus-AGP21 expression. Plasmolysis was done using 800 mM mannitol.

### Molecular dynamics (MD) simulations

MD simulations were performed on two non-glycosylated and seven glycosylated Ala1-Pro2-Ala3-Pro4-Ser5-Pro6-Thr7-Ser8 (APAPSPTS) peptides, in which the starting structure was constructed as a type-II polyproline helix, with ϕ ~ −75 and ψ ~ 145. The non-glycosylated motifs differ by the presence of alanine (AAAASATS), proline (APAPSPTS) or 4-*trans*-hydroxyproline (AOAOSOTS) residues. At the same time, the glycosylated motifs reflect different peptide glycoforms, constructed as full glycosylated (AOAOSOTS). Every *O*-glycosylation site was filled with an arabinogalactan oligosaccharide moiety (**Supplementary Item 5**), in which the *O*-glycan chains and carbohydrate-amino acid connections were constructed based on the most prevalent geometries obtained from solution MD simulations of their respective disaccharides, as previously described (Pol-Fachin & Verli 2012), thus generating the initial coordinates for glycopeptide MD calculations. Such structures were then solvated in rectangular boxes using periodic boundary conditions and the SPC water model (Berendsen et al. 1984). Both carbohydrate and peptide moieties were described under GROMOS96 43a1 force field parameters, and all MD simulations and analyses were performed with GROMACS simulation suite, version 4.5.4 (Hess et al. 2008). The Lincs method (Hess et al. 1997) was applied to constrain covalent bond lengths, allowing an integration step of 2 fs after an initial energy minimization using the Steepest Descents algorithm. Electrostatic interactions were calculated with the generalized reaction-field method Tironi et al. (1995). Temperature and pressure were kept constant at 310 K and 1.0 atom, respectively, by coupling (glyco)peptides and solvent to external baths under V-rescale thermostat Bussi et al. 2007) and Berendsen barostat (Berendsen et al. 1987) with coupling constants of *t* =0.1 and *t*=0.5, respectively, via isotropic coordinate scaling. The systems were heated slowly from 50 to 310 K, in steps of 5 ps, each one increasing the reference temperature by 50 K. After this thermalization, all simulations were further extended to 100 ns. See **Table S3**.

## Acknowledgements

We thank ABRC (Ohio State University) for providing T-DNA lines seed lines. We would like to thank Dr. Sigal Savaldi-Goldstein for providing GL2p∷BRI1-GFP, COBL9p∷BRI1-GFP and Gl2p∷BRI1-GFP + COBL9p∷BRI1-GFP reporter lines, Dr. Paul Dupree for providing *fut4* and *fut6* mutant lines, Dr. Santiago Mora García for providing *bzr1* and *bes1* mutant lines, Dr. Ana Caño Delgado for providing seeds of BRI1-GFP, *bin2-1* and BZR-YFP, and Dr. Gustavo Gudesblat for providing *gsk triple* mutant seeds. Dr. Malcolm Bennet and Dr. Liam Dolan for the RHD6-GFP and RSL4-GFP lines. This work was supported by grants from ANPCyT (PICT2014-0504, PICT2016-0132 and PICT2017-0066) and ICGEB CRP/ARG16-03 to J.M.E.

## Author Contribution

C.B, J.G.D and M.M.R performed most of the experiments, analysed the data and wrote the paper. L.P.F and H.V. performed molecular dynamics simulations and analysed this data. M.C.S analysed the phenotype of glycosyltransferase mutants and BRI1-GFP reporters. B.V. analysed the molecular dynamics simulations data. M.C synthesized α-Man-Y and β-Glc-Y reagents. G.S. commented on the project, read the manuscript, and commented on the results. S.M. and E.M. analysed the data and commented on the results. J.M.P., D.R.R.M., Y.R., and S.M.V commented on the results. J.M.E. designed research, supervised the project, and wrote the paper. This manuscript has not been published and is not under consideration for publication elsewhere. All the authors have read the manuscript and have approved this submission.

