## Supplementary Information for "A cell surface arabinogalactan-peptide influences root hair cell fate"

**Supplemental Information**

Supplementary Figures S1-S7

Supplementary Tables S1-S4

Supplementary References

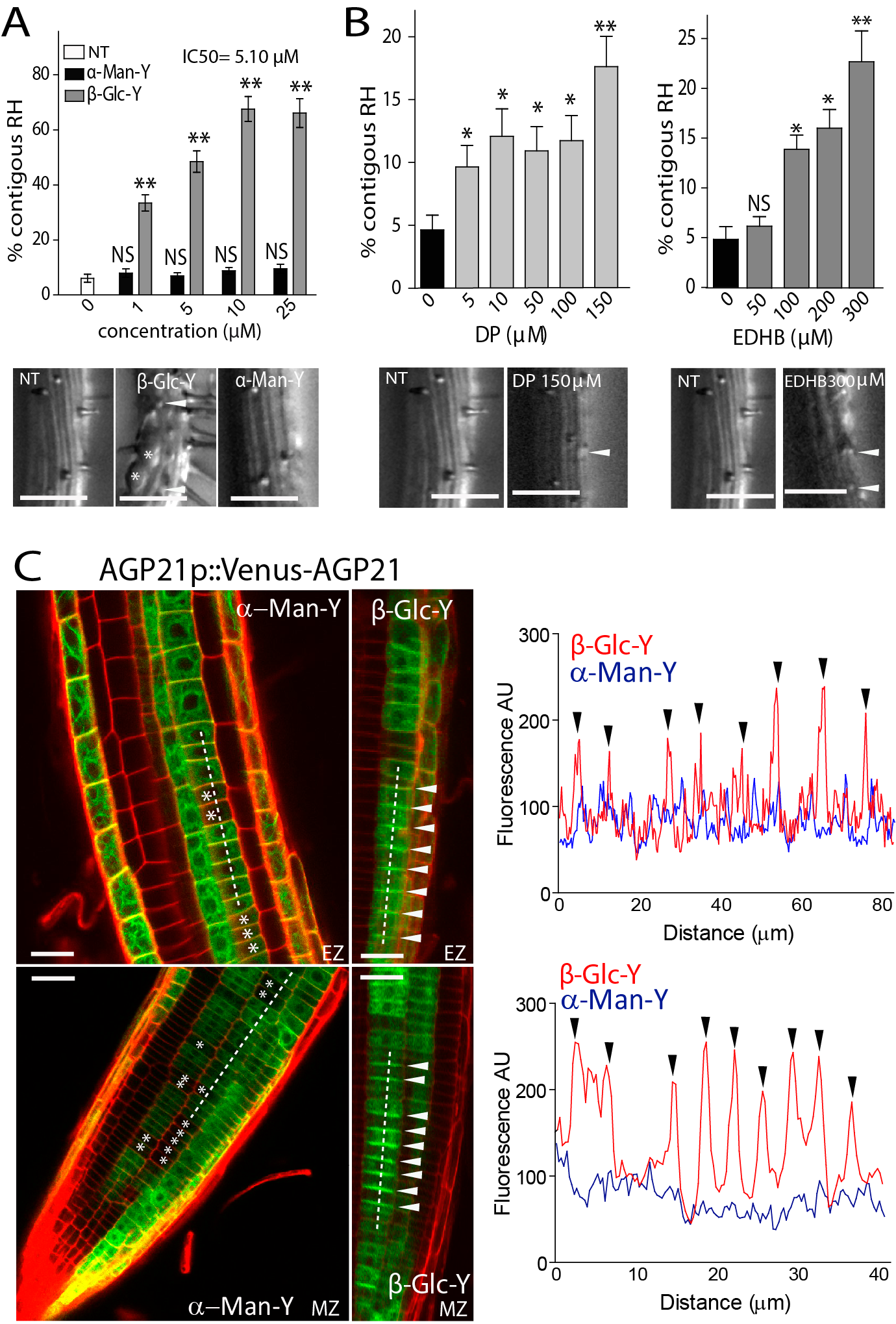

**Figure S1. Perturbation of *O*-glycosylated AGPs affect RH cell fate program**. **β-Glc-Y is able to trigger an over-accumulation of AGP21 peptide in the cell surface.**

(A) Contiguous RH phenotype developed under pharmacological of AGPs. Pictures below each graph indicate the RH phenotype in detail. Arrowheads indicate two contiguous RH cell protuberances. Asterisk indicated bulging cells. Roots treated with β-Glucosyl Yariv (β-Glc-Y) or with α-Mannosyl Yariv (α-Man-Y) as control. NT= non-treated.

(B) Contiguous RH phenotype developed under pharmacological disruption of peptidyl-proline. Roots treated with two distinct P4H inhibitors, α,α-dipyridyl (DP) and ethyl-3,4-dihydroxybenzoate (EDHB). Pictures below each graph indicate the RH phenotype. Arrowheads indicate two contiguous RH cell protuberances. Asterisk indicated bulging cells. Scale bar= 50 μm.

(C) AGP21 is expressed in some but not all trichoblast and atrichobast cells with a discontinuous pattern (asterisk) in the meristematic zone (MZ) and elongation root zones (EZ). Some root epidermal cell layer lack AGP21 (line). On the left, the effect of β-Glucosyl Yariv (β-Glc-Y) on the accumulation of AGP21p::V-AGP21 on root epidermal cells. Arrowheads indicate cell surface AGP21 peptide over-accumulation in transversal walls in the treated roots with β-Glc-Y. Scale bars= 10 μm. Plot profiles (dashed lines) indicates the accumulation of AGP21p::V-AGP21 (black arrowheads) when roots are treated with β-Glc-Y.

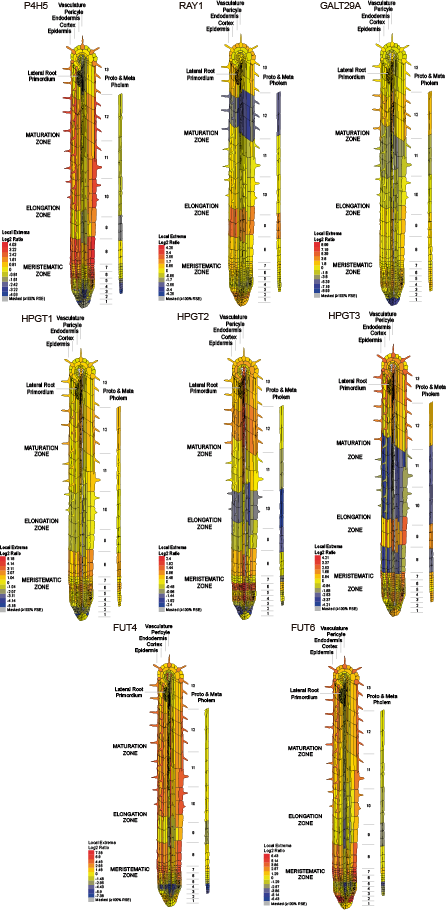

**Figure S2. Expression pattern of enzymes involved in proline hydroxylation and *O*-glycosylation of AGPs in the *Arabidopsis* roots.**

Expression of P4H5, GALT29A, RAY1, HPTG1-HPGT3 and FUT4/FUT6 are based on ePlant server (http://bar.utoronto.ca/eplant/), tissue and experiments eFP viewers. Most of these AGP-modifying enzymes are highly expressed in epidermal cells.

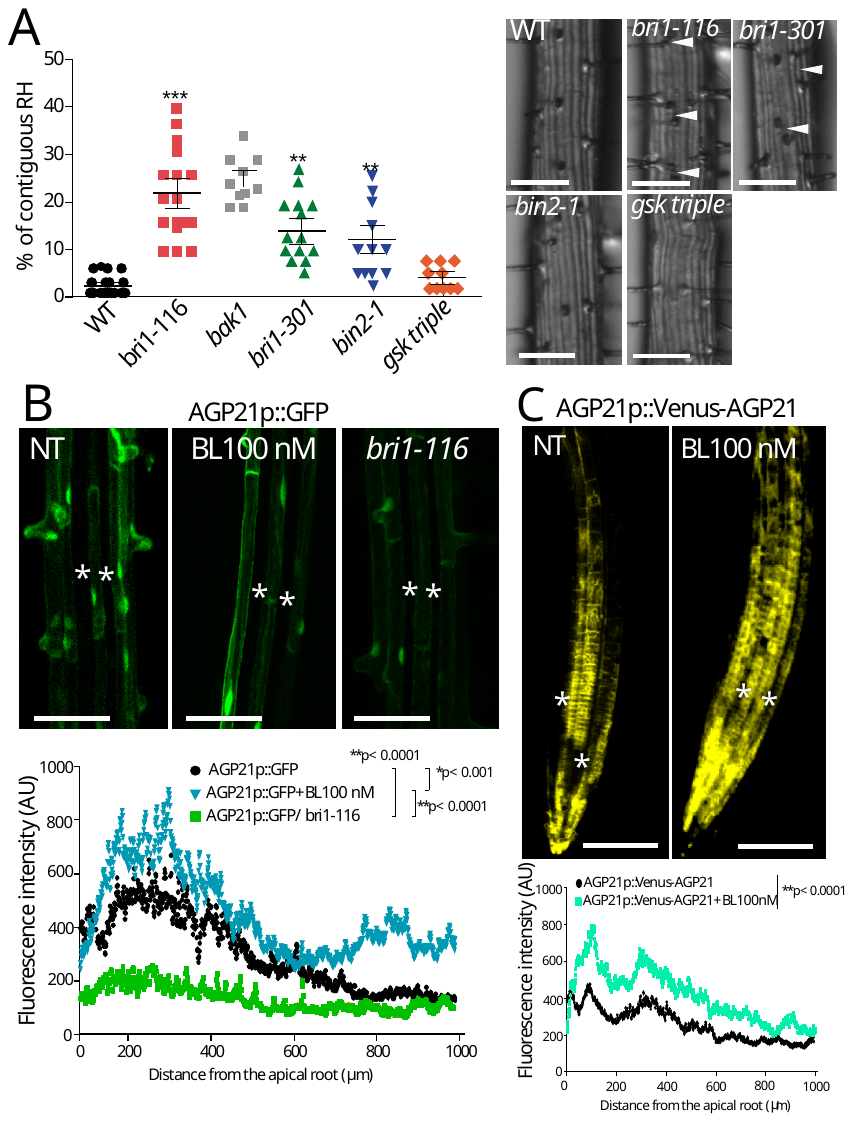

**Figure S3. BR deficiency triggers RH abnormal development and BR control of AGP21 expression.**

(A) Contiguous RH phenotype developed in BR constitutive active *bin2-1* and *gsk* *triple* (mutant mutant *bin2-3,bil1,bil2*) as well as in *bri1* mutants. Pictures indicate the RH phenotype in detail. Arrowheads indicate two contiguous RH cell protuberances. Scale bar= 50 μm. *P*-value of one-way ANOVA, (***) P<0.0001, (**) P<0.001.

(B) Effect of 100 mM BL (brassinolide, an active form of BR) or *bri1-116* mutation on the expression of AGP21p::GFP transcriptional reporter. Quantification of V-AGP21 intensity signal is indicated along the root axis in each treatment. Scale bar= 50 μm. Asterisk indicates lack of expression in atrichobast cells. Scale bar= 50 μm.

(C) Effect of BL on the expression of the AGP21p::V-AGP21 protein reporter. Quantification of V-AGP21 intensity signal is indicated along the root axis in each treatment. Scale bar= 200 μm. Asterisk indicates lack of expression in atrichobast cells.

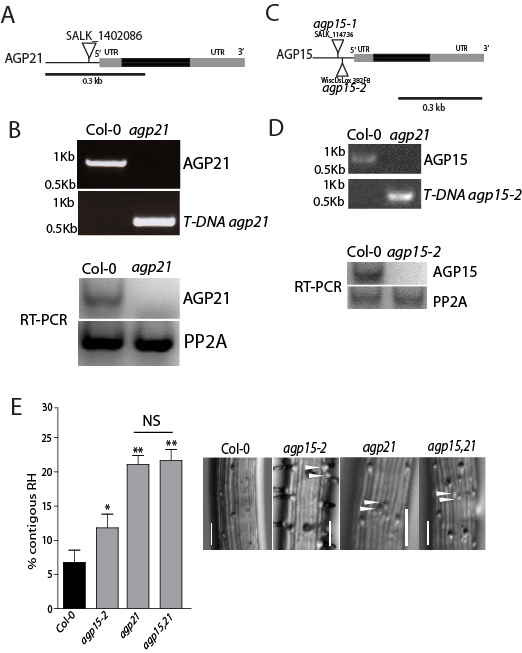

**Figure S4. *agp21* and *agp15* mutants characterization.**

(A) Schematic representation of AGP21 peptide. Position of *agp21* T-DNA insertion is indicated in the promoter region**.**

(B) Validation of *agp21* T-DNA mutant line. Total RNA was extracted from 10 days old roots. PP2A was used as control. The primers used for RT-PCR are listed in **Table S3**.

(C) Schematic representation of AGP15 peptide. Positions of T-DNA insertions are indicated. Prom: promoter region**.**

(D) Validation of *agp15* T-DNA mutant lines. Total RNA was extracted from 10 days old roots. PP2A was used as a control. The primers used for RT-PCR are listed in **Table S3**.

(E) Contiguous RH phenotype developed in the genetic disruption of multiple AGP peptides. Pictures below each graph indicate the RH phenotype in detail. *P*-value of one-way ANOVA, (**) P<0.001, (*) P<0.01. NS= not significant different. Error bars indicate ±SD from 3 biological replicates. Arrowheads indicate two contiguous RH cell protuberances. Scale bar= 50 μm.

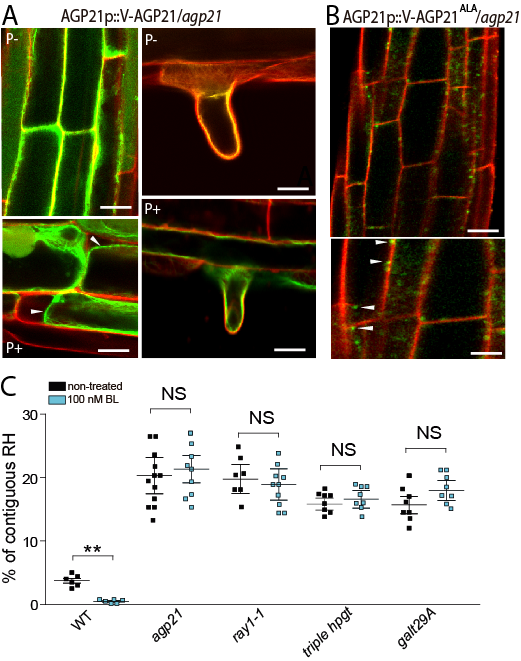

**Figure S5. AGP21 expression at the cell surface in epidermis and RH cells**. **Perception of BR in epidermal cells is abolished in the *agp21* and related under-*O*-glycosylated AGP mutants**.

(A) Expression of AGP21p::V-AGP21 at the plasma membrane in non-plasmolyzed (P-) and plasmolyzed (P+) epidermal cells (on the right) and RHs (on the left) with 800 mM Mannitol. Arrowheads indicate retraction of plasma membrane. Scale bar= 10 μm.

(B) Expression of AGP21p::V-AGP21^ALA^. This version of AGP21 peptide accumulates as intracellular dots. Scale bar= 10 μm.

(C) Contiguous RH phenotype in WT Col-0, *agp21*, *ray1-1*, *triple hpgt* and *galt29A* in non-treated and treated roots with 100nM BL. *P*-value of one-way ANOVA, (**) P<0.001. NS= not significant different. Error bars indicate ±SD from biological replicates.

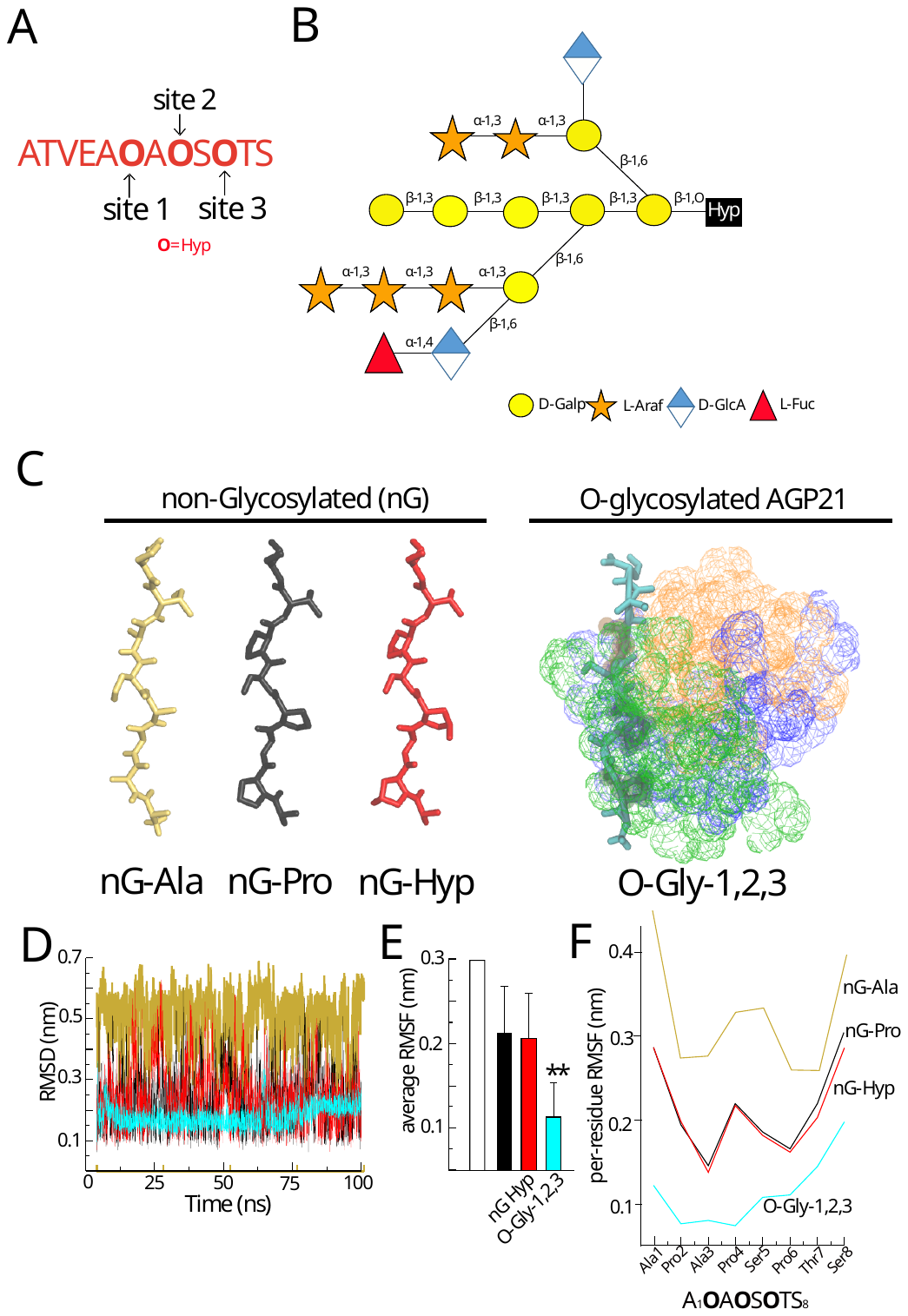

**Figure S6**. ***O*-glycans provide stability to the AGP21 peptide conformation.**

(A) Three putative *O*-glycosylation sites present in the mature AGP21 peptide ATVEAOAOSOTS (O=Hyp).

(B) Arabinogalactan oligosaccharide (AG) composed of 15 residues including L-Glucuronic Acid (GlcA) and L-Fucose attached to each Hyp units used during MD simulations. Schematic representation of the arabinogalactan oligosaccharide used to construct AOAOSOTS peptide glycosylated form. Mutants for some GTs used in this study (*p4h5*, triple *hpgt*, *galt29A*, *ray1* and *fut4 fut6*) are indicated.

(C) Most representative structure for the simulated glyco-peptides, in which the protein moiety is shown as sticks (with *O*-glycosylated amino acids as VDW) and the carbohydrate moieties are shown as dots. The *O*-glycan chains linked to site 1 (Hyp_2_) are colored as green; to site 2 (Hyp_4_) as blue; and to site 3 (Hyp_6_) as orange. *O*-Gly-1,2,3 refers to a fully *O*-glycosylated AGP21 peptide.

(D) All-atom Root Mean Square Deviation (RMSD) for APAPSPTS protein moiety in the performed MD calculations.

(E) Root Mean Square Fluctuation (RMSF) obtained by averaging the per-residue values in each peptide simulation. nG abbreviates non-glycosylated; *O*-Gly-1,2,3 refers to a fully *O*-glycosylated AGP21 peptide.

(F) Per-residue RMSF for each MD condition.

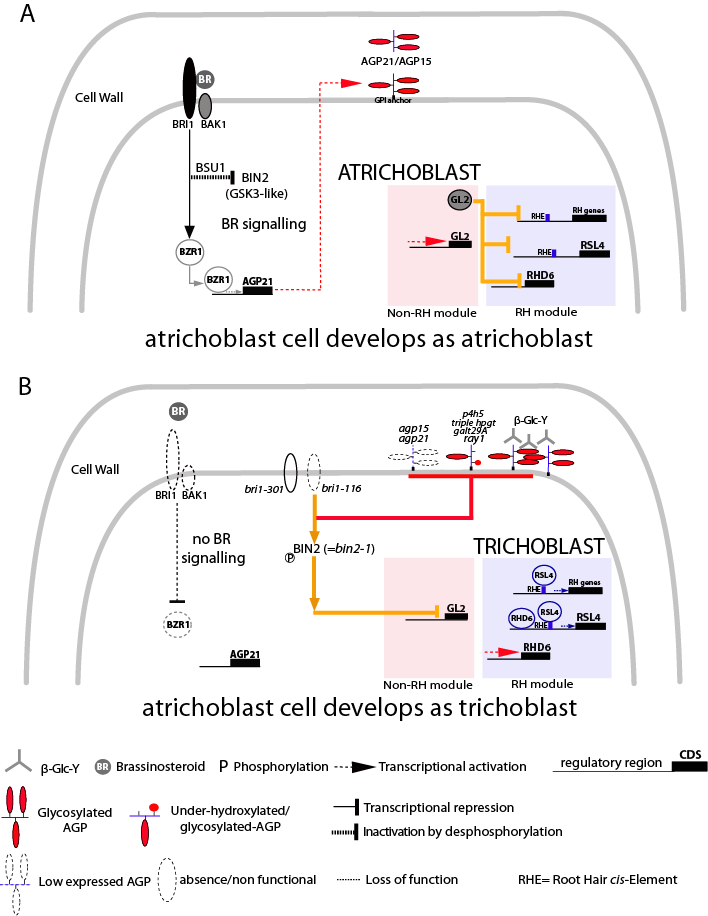

**Figure S7. AGP21 peptide influences RH cell fate in a BIN2-dependent manner.**

(A) Under normal BR-signaling. Model of BR-BRI1-BAK1 pathway modulation of RH cell fate and the plasma membrane *O*-glycosylated AGP21. BR-signalling controls RH cell fate by inhibiting phosphorylation activity of BIN2 and impacting on GL2 expression. Most atrichoblast cells keep their cell fate identity.

(B) Disruption of *O*-glycosylated AGPs (with β-Glc-Y) and the lack of AGP21 (in *agp21* mutant) as well as abnormal glycosylated AGPs (in *p4h5, triple hpgt, ray1, galt29A* mutants and DP/EDHB treated roots) may interfere with BRI1-brassinosteroid responses and BIN2 downstream effect on the RH repressor GL2. Some atrichoblast cells lost their cell fate identity and they develop ectopic RH cells. As a consequence, GL2 transcriptional repression triggers RH development in atrichoblast cells producing contiguous RH. In this case the atricoblast cell differentiates as trichoblast. In addition, disruption of AGP21 peptide (and AGPs) act in a BR-independent manner but converges on BIN2 to triggers abnormal RH cell fate.

**Table S1**. GTs involved in AGP modification used in this study.

| **Protein/Gene code** | **CAZY family** | **Full name/ Activity** | **References** |
| --- | --- | --- | --- |
| **Enzymes acting on AGP and related glycoproteins** | | | |
| **HPGT1/**AT5G53340  **HPGT2/**AT4G32120  **HPGT3/**AT2G25300 | GT31 | Hydroxyproline Galactosyltransferases 1-3/  transfer of D-Gal*p* to the hydroxyl-group on Hyp residues in AGPs and related glycoproteins | Egelund et al. 2007 |
| **GalT29A/**AT1G08280 | GT29 | Galactosyltransferase 29A/  transfers galactose in β-D-(1→6) position as elongating enzyme or in β-D-(1→3)-galactosyl oligosaccharides | Dilokpimol et al. 2014; Geshi et al. 2013 |
| **RAY1/**AT1G70630 | GT77 | Reduced Arabinose Yariv 1/  β-arabinofuranosyltransferase that adds arabinose on the AGP backbone | Gille et al. 2013 |
| **FUT4/**AT2G15390  **FUT6/**AT1G14080 | GT77 | Fucosyltransferase/  transfers fucose as α-L-(1→2) position to α-L-Ara*f*-(1→) unit | Tryfona et al 2014 |
| **Enzymes acting on EXTs and related glycoproteins** | | | |
| **RRA3**/AT1G19360 | GT77 | Reduced Residual Arabinose 3/ Arabinosyltransferase that transfers of β-L-Ara*f* -(1→2) units | Egelund et al. 2007  Velasquez et al. 2011 |
| **SGT1 (SERGALT1)** /At3g01720 | GT96 | peptidyl serine *O*-α-galactosyltransferase | Saito et al. 2014 |

**Table S2**. Mutants and transgenic lines generated and used in this study.

| Gene name | AGI code | Genetic background | Transgenic line | Reference |
| --- | --- | --- | --- | --- |
| RRA3 | AT1G19360 | *rra3* | - | Egelund et al. 2007 Velasquez et al. 2011 |
| SGT1 | At3g01720 | *sgt1 (sergalt1) rra3* | - | Saito et al. 2014  Velasquez et al. 2015a |
| P4H5 | AT2G17720 | *p4h5* | - | Velasquez et al. 2011 |
| GalT29A | AT1G08280 | *galt29A* | - | Dilokpimol et al. 2014; Geshi et al. 2013 |
| RAY1 | AT1G70630 | *ray1* | - | Gille et al. 2013 |
| AGP15 | AT5G11740 | *agp15*  *agp15,agp21* | - | This work  This work |
| **AGP21** | AT1G55330 | *agp21*  *agp21*  *agp21*  Col-0  Col-0 | -  35Sp::Venus-AGP21  AGP21p::Venus-AGP21  35Sp::Venus-AGP21  AGP21p::GFP | This work  This work  This work  This work  This work |
| **Brassinosteroid lines** | | | | |
| **BRI1** | AT4G39400 | *bri1-5*  *bri1-116*  *bri1-116* | -  -  AGP21p::GFP  35Sp::BRI1-GFP | Noguchi et al. 1999  Li & Chory 1999  This work |
| **BZR1** | AT1G75080 | bzr1-D  Col-0 | BZR1p::BZR1-YFP  35Sp::BZR1-GFP  BZR1-D  *bzr1-1 crispr-cas* | Wang et al. 2002  Chaiwanon et al. 2015  Saito et al. 2018 |
| **BES1** | AT1G19350 | bes1-D | -  35Sp::BES1-GFP  BES1-D  *Bes1-1 bzr1-1 crispr-cas* | Yin et al. 2002  Saito et al. 2018 |
| **BIN2** | AT4G18710 | *bin2-1*  *bin2,bil1,bil2* | BIN2p:BIN2-GFP | Yin et al. 2002 |
| **BIL1** | AT2G30980 |  | - | Kim 2012  Yan et al. 2009 |
| **BIL2** | AT1G06390 |  | - |  |
| **Trichoblast and Atrichoblast marker lines** | | | | |
| **RHD6** | AT1G66470 | Wt Col-0 | RHD6p::RHD6-GFP | Yi et al. 2010 |
| **RSL4** | AT1G27740 | Wt Col-0 | RSL4p::RSL4-GFP | Yi et al. 2010 |
| **GL2** | AT1G79840 | Wt Col-0 | GL2p::GL2-GFP | Lin et al. 2015 |
| **EXP7** | AT1G12560 | Wt Col-0 | EXP7p::nGFP | Kim et al. 2006 |

**Table S3**. Average ζ angle values* during the performed MD simulations of AGP21 peptide.

| **Peptide state** | **ζ angle 1**  **Ala1-Pro2-Ala3-Pro4** | | **ζ angle 2**  **Pro2-Ala3-Pro4-Ser5** | | **ζ angle 3**  **Ala3-Pro4-Ser5-Pro6** | | **ζ angle 4**  **Pro4-Ser5-Pro6-Thr7** | | **ζ angle 5**  **Ser5-Pro6-Thr7-Ser8** | |
| --- | --- | --- | --- | --- | --- | --- | --- | --- | --- | --- |
| Type-II polyproline | | -110 ± 15 | | -110 ± 15 | | -110 ± 15 | | -110 ± 15 | | -110 ± 15 |
| non-Glyco, Ala AGP21 | | **159 ± 89** | | **149 ± 116** | | **153 ± 126** | | **166 ± 97** | | **-177 ± 75** |
| non-Glyco, Pro AGP21 | | -124 ± 46 | | -125 ± 31 | | -130 ± 49 | | -130 ± 31 | | -149 ± 49 |
| non-Glyco, Hyp  AGP21 | | -130 ± 47 | | -125 ± 31 | | -133 ± 46 | | -128 ± 32 | | -150 ± 51 |
| Glyco1,2,3  AGP21 | | **-112 ± 25** | | **-175 ± 16** | | **-168 ± 22** | | **-150 ± 21** | | **-131 ± 43** |

* Average ± standard deviation values measured for the second half of MD simulations (from 50 ns to 100 ns).

**Table S4**. Primers used in this study.

| **Purpose** | **Name** | **Sequence (5’ to 3’)** |
| --- | --- | --- |
| **RT-PCR AGP15** | Forward | CATCGGCACAATCTGAGG |
|  | Reverse | ACCATCACAGTAACTTAGATCC |
| **RT-PCR AGP21** | Forward | GCAATGAAGATGAAGATGATGG |
|  | Reverse | TCAGAAGTTGGGCTTGGAG |
| **RT-PCR PP2A** | Forward | TCCGAGATCACATGTTCCAAACTC |
|  | Reverse | CCGTATCATGTTCTCCACAACCG |
| **T-DNA AGP15** | Forward | GACACGAAAGACGCTGAGATC |
|  | Reverse | AGGAGAAATTTGCACCCATTC |
| **T-DNA AGP21** | Forward | TTTGGTGTGAACGTTGGTATG |
|  | Reverse | CAAAAGATGAAACCAGATGCC |
| **T-DNA GALT29A** | Forward | TTTGTGGCTCGAGTAAACCC |
|  | Reverse | AAGCATGAGATTGTGATTCGG |
| **T-DNA RAY1** | Forward | TTTGGAGCGTATGGATCAAAG |
|  | Reverse | GAGTTATGCTCACGAGCTTGG |
| **Promoter AGP21** | Forward | TAATGCCAACTTTGTACAAAAAAGCAGGCT |
|  | Reverse | CCCAGCTTTCTTGTAC |
| **Venus-AGP21** | Forward | GGGGACAAGTTTGTACAAAAAAGCAGGCTTAACCatggaggcaatgaagatg |
|  | Reverse | GGGGACCACTTTGTACAAGAAAGCTGGGTCtcaaaagatgaaaccaga |
| **RT-PCR AGP15** | Forward | CATCGGCACAATCTGAGG |
|  | Reverse | ACCATCACAGTAACTTAGATCC |
| **RT-PCR AGP21** | Forward | GCAATGAAGATGAAGATGATGG |
|  | Reverse | TCAGAAGTTGGGCTTGGAG |

### Geshi, N., Johansen, J.N., Dilokpimol, A., Rolland, A., Belcram, K., Verger, S., et al. (2013). A galactosyltransferase acting on arabinogalactan protein glycans is essential for embryo development in Arabidopsis. *Plant J.* 76, 128–137.doi: 10.1111/tpj.12281

### [Gille S](https://www.ncbi.nlm.nih.gov/pubmed/?term=Gille%20S%5BAuthor%5D&cauthor=true&cauthor_uid=23396039), [Sharma V](https://www.ncbi.nlm.nih.gov/pubmed/?term=Sharma%20V%5BAuthor%5D&cauthor=true&cauthor_uid=23396039), [Baidoo EE](https://www.ncbi.nlm.nih.gov/pubmed/?term=Baidoo%20EE%5BAuthor%5D&cauthor=true&cauthor_uid=23396039), [Keasling JD](https://www.ncbi.nlm.nih.gov/pubmed/?term=Keasling%20JD%5BAuthor%5D&cauthor=true&cauthor_uid=23396039), [Scheller HV](https://www.ncbi.nlm.nih.gov/pubmed/?term=Scheller%20HV%5BAuthor%5D&cauthor=true&cauthor_uid=23396039), [Pauly M](https://www.ncbi.nlm.nih.gov/pubmed/?term=Pauly%20M%5BAuthor%5D&cauthor=true&cauthor_uid=23396039). (2013). Arabinosylation of a Yariv-precipitable cell wall polymer impacts plant growth as exemplified by the Arabidopsis glycosyltransferase mutant ray1. [*Mol Plant*.](https://www.ncbi.nlm.nih.gov/pubmed/23396039) 6(4):1369-1372. doi: 10.1093/mp/sst029

Hess B, Kutzner C, Van Der Spoel D, Lindahl E (2008) GROMACS 4: Algorithms for highly efficient, load- balanced, and scalable molecular simulation. *Journal of Chemical Theory and Computation* 4:435-444

Hess B, Bekker H, Berendsen H.J.C., Fraaije J.G.E.M. (1997) LINCS: A linear constraint solver for molecular simulations. *Journal of Computational Chemistry* 18:1463-1472

Houbaert A et al. (2018) POLAR-guided signaling complex assembly and localization drive asymmetric cell division. *Nature* 563(7732), 574-578

Kim T-W, et al. (2012). Brassinosteroid regulates stomatal development by GSK3-mediated inhibition of a MAPK pathway. *Nature* 482(7385), 419–422. <http://doi.org/10.1038/nature10794>

Kim, D.W. et al. (2006). Functional Conservation of a Root Hair Cell-Specific cis-Element in Angiosperms with Different Root Hair Distribution Patterns. *Plant Cell* 18(11), 2958–2970

Noguchi, T., Fujioka, S., Choe, S., Takatsuto, S., Yoshida, S., Yuan, H., … Tax, F. E. (1999). Brassinosteroid-Insensitive Dwarf Mutants of Arabidopsis Accumulate Brassinosteroids. *Plant Physiology* 121(3), 743–752

Pol-Fachin L, Verli H (2012) Structural glycobiology of the major allergen of Artemisia vulgaris pollen, Art v 1: O-glycosylation influence on the protein dynamics and allergenicity. *Glycobiology* 22:817-825

Saito, F., Suyama, A., Oka, T., Yoko-o, T., Matsuoka, K., Jigami, Y., and Shimma, Y. (2014). Identification of novel peptidyl serine O-galactosyltransferase gene family in plants. *J. Biol. Chem.* 30:20405–20420

Saito M., Kondo Y., and Fukuda H. (2018) BES1 and BZR1 redundantly promote phloem and xylem differentiation. *Plant Cell Physiol.* 2018 59(3):590-600

Tironi IG, Sperb R, Smith PE, van Gunsteren WF (1995) A generalized reaction field method for molecular-dynamics simulations. *The Journal of Chemical Physics* 102:5451-5459

Wang Z-Y, Nakano T, Gendron J, He J, Chen M, Vafeados D, Yang Y, Fujioka S, Yoshida S, Asami T & Chory J (2002). Nuclear-Localized BZR1 Mediates Brassinosteroid-Induced Growth and Feedback Suppression of Brassinosteroid Biosynthesis. *Developmental Cell* 2, 505–513. doi.org/10.1016/S1534-5807(02)00153-3

Yan Z, Zhao J, Peng P, Chihara RK, Li J. (2009). BIN2 functions redundantly with other Arabidopsis GSK3-like kinases to regulate brassinosteroid signaling. *Plant Physiology* 150:710–721. doi: 10.1104/pp.109.138099

Yin Y, Wang ZY, Mora-Garcia S, Li J, Yoshida S, Asami T, Chory J. (2002) BES1 accumulates in the nucleus in response to brassinosteroids to regulate gene expression and promote stem elongation. *Cell* 109(2):181-91
